# Removing mains power artefacts from EEG – a novel template-based method

**DOI:** 10.1101/2020.01.27.911586

**Authors:** Shenghuan Zhang, Brendan McCane, Phoebe S-H Neo, Neil McNaughton

## Abstract

EEG signals are often contaminated with artefacts, particularly with mains power from electrical equipment. Low-pass filtering and notch filtering can lose valuable data. Here we describe a novel mains power noise removal method based on the fact that mains power noise is a sine wave component with an essentially fixed frequency and the same phase across all channels. This removes the mains power component, leaving uncontaminated EEG largely unchanged. Mains removal had a success rate of >99.9% recovered variance of the original EEG when removing synthesised mains power components. We compared this method with two other popular mains power noise removal methods. Our method was better than Cleanline which is an ICA based method and multiple PLI which is an adaptive notch filter based method. With the higher recovery, our method shows clear advantages over Cleanline and multiple PLI for removing mains power noise. From the aspect of data loss, it is obviously better than low-pass filter and normal notch filter.

## 1. Introduction

The complex waveforms of the EEG contain much valuable information. However, artefacts may interfere with the signals that we are interested in. Physiological sources contaminate the brain’s EEG with signals generated by activity in other organs and tissues, such as eye blinks, eye movement, cardiac activity and other muscle movement, while non-physiological artefacts are generated by changes in impedance between the electrodes and the scalp, various types of electromagnetic signal in the environment and, particularly, the electromagnetic signal coming from mains power (Uriguen & Garcia-Zapirain, 2015). Mains power generally produces a substantial local peak in the EEG power spectrum around the frequency of electrical power supply (50 Hz in some countries and 60 Hz in others).

In some EEG research, frequencies over 30 Hz are filtered using a low-pass filter. In this situation, the 50 or 60 Hz noise will not have a major influence on the research results. However, high frequency band oscillations (particularly in the gamma range) are important in some studies especially of emotion, mental health and working memory (Howard, 2003; Li & Lu, 2009; Nugent et al., 2018). Simple filtering of this band of EEG signal will lead to unacceptable data loss.

Several methods have been designed to reduce the influence of mains power noise on EEG signals of similar frequency but each has its limitations. The most common (and arguably the most important) method is to use an electrical ground wire and shielding to restrain the power frequency interference. Appropriate choice of a recording reference electrode is also important. However, after applying these hardware level methods, there is usually a large residual mains interference component in the signal (Thorp & Steinmetz, 2008). Therefore, signal processing methods at the software level have been designed to further remove mains power noise. A widely used method is the notch filter which rejects the signal in the close vicinity of the mains power frequency (~50 or ~60 Hz). However, notch filtering generally creates band-holes at the target frequency, with the same problem as the low-pass filter that important EEG data are lost. A recent method, which has been especially widely used with EEG data, is to “estimate and remove sinusoidal artefacts from ICA components or scalp channels using a frequency-domain (multi-taper) regression technique with a Thompson F-statistic for identifying significant sinusoidal artefacts” (Mitra, 2007; Mullen, 2012). This method is called CleanLine and is available as a function within the EEGLAB plug-in of MATLAB (Brunner et al., 2013; Delorme and Makeig, 2004). This method uses a sliding window to adaptively estimate sine wave amplitude to subtract. It does not need an extra electrode to record noise and it does not create band-holes. Another robust and widely used method of mains power removal from neural recordings is multichannel PLI (Keshtkaran, 2014). Like CleanLine, this method does not require a reference signal and does not create band-holes. It uses an adaptive notch filter to estimate the fundamental frequency and harmonics of the mains power noise. Then a modified recursive least squares algorithm is used to estimate the amplitude and phase of each harmonic.

Here, we propose an approach, similar to CleanLine, for mains power noise removal that is based on the assumption that the phase of the mains power signal is constant across all channels. It obtains the noise signal by summing signals across all channels *for the entire recording* to reduce the contribution of real EEG and strengthen the contribution of mains power to the noise signal. Then the phase of the noise is then calculated based on the resultant noise-enhanced signal. A unit amplitude sinewave is then created using the observed phase and fixed frequency value. Finally, the unit sinewave is fitted back to each channel and the amplitudes of the noise signal of each channel are obtained. In the results, we compare this method with CleanLine and multichannel PLI applied to the same data.

## 2. Method and Materials

### 2.1 Participants

Determination of mains power parameters of the key population and of the algorithm’s effectiveness were carried out on data obtained from 73 right-handed participants (44 females, 29 males; aged 18-37 years with a mean of 21.56 years). These data were obtained as part of a separate experimental study. All procedures were approved by the University of Otago Ethics Committee (approval number: H15/005). All participants were recruited through the University of Otago Student Job Search, provided informed consent before participating in the experiment, and received NZ$15 per hour as compensation for the time and effort spent in attending the experiment.

### 2.2 Procedure

Prior to EEG testing, the experimenter measured each participant’s head circumference and marked Fp1 and Fp2 according to the International 10-20 system (Sharbrough, 1991) using a black marker. The participant was then fitted with a Waveguard (Ag/Agcl) EEG cap with Gnd as the ground electrode and M1 and M2 (mastoids) recorded separately and then averaged as reference. The cap was connected to an ANT system (Advanced Neuro Technology B.V., Enschede, The Netherlands). Electrode gel (Electro-Cap International, Eaton, OH, USA) was inserted into 20 electrodes via a blunt square-tipped 16-gauge needle (Precision Glide, Needle, Becton Dickinson, Franklin Lakes, NJ, USA); impedance reduced to <5KΩ by gentle abrasion of the scalp with the tip of the needle; and brief relaxation-induced alpha rhythm and deliberate eye-blink traces assessed by the experimenter to ensure good recording, with adjustments as necessary (preparation time ~ 30 minutes).

A relaxation test was then performed by instructing the participants to remain relaxed, with their eyes open and then closed for one-minute intervals, the order was O-C-O-C-C-O-C-O (C: eyes closed, O: eyes open). Resting EEG was recorded throughout this period (8 minutes duration).

The data analysed were from 18 primary recorded channels: Fp1, Fpz, Fp2, F7, F3, Fz, F4, F8, T7, C3, Cz, C4, T8, P7, P3, Pz, P4 and P8, which were referenced to CPz when recording, and then re-referenced to the average of ‘A1’ and ‘A2’ for analysis. The signals were then filtered by 1-Hz high-pass filter to remove baseline drift. The sampling rate was 256Hz.

### 2.3 Algorithm implementations

Our custom algorithms were implemented with Matlab R2018a (9.4.0.813654). Function ‘robustfit (X,y)’ was used to return the coefficient estimates of X and y using robust regression (Dumouchel & O’Brien, 1989; Holland, Welsch, & Methods, 1977). For the purposes of comparison with other mains power noise removal procedures, we implemented the CleanLine method using the function, pop_CleanLine (), in EEGLAB (v13.4.4). This function was implemented with Matlab R2016 (Matlab R2018a tended to crash when running EEGLAB). The source code of the multichannel PLI method was downloaded from open access at https://github.com/mrezak/removePLI. It was implemented with Matlab R2018a.

### 2.4 Mains power removal method

We assumed that mains power should have a fixed phase all through EEG recordings and be synchronous across all channels whereas brain signals are not the same phase as each other and should not be synchronous. Therefore, we designed a phase-fixed method to remove the mains noise component in EEG signal.

The mains power method consists of four key steps. The first identifies the frequencies at which the mains power occur. The second averages signals across all channels to reduce the real brain signals relative to the mains noise. Next, an FFT algorithm was used to calculate the phase of mains power at the appropriate frequency (see below). The phase and the predetermined frequency values were then used to create a unit-amplitude sine wave as a noise template. Finally, the fitted unit signal was scaled to each channel separately with the robust regression technique (Dumouchel & O’Brien, 1989) and the result was subtracted to leave residual EEG. The whole procedure was then repeated for the next frequency (see below).

#### 2.4.1 Determining the frequency range of the mains noise

According to the electricity (safety) regulations (New Zealand Parliamentary Counsel Office, 2010), the frequency of electricity supplied must be maintained between 47 Hz and 53 Hz. Figure 1 shows an example of an EEG signal which was contaminated by the mains power signal in our dataset. The signal, transformed into the frequency domain by FFT, shows a peak from 49.9 Hz to 50.1 Hz. Visual inspection of all individual’s FFT spectrum confirmed that all mains power signal in our dataset falls into this range.

**Figure 1.**
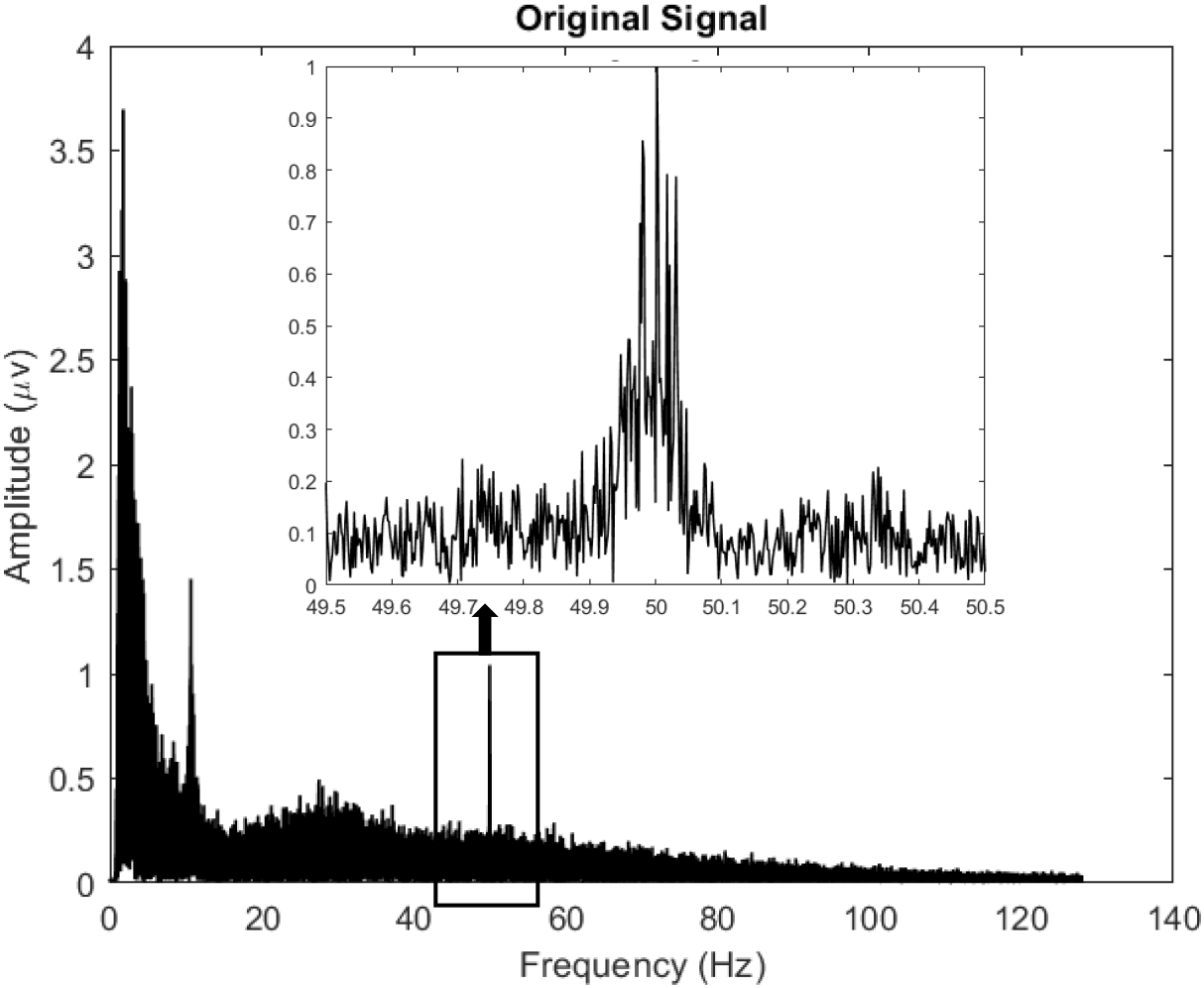
FFT spectrum before mains noise removal. The spectrum of a single-channel EEG signal from a randomly selected participant clearly shows that there is mains power interference in the range from 49.9 Hz to 50.1 Hz.

#### 2.4.2 Averaging signals of all channels to enhance the mains noise

On the assumption that across all recording channels, EEG should show different, and mains noise signal should show the same phases, we average the signals across all the channels to reduce the random phase EEG component and extract the fixed phase noise component. Mathematically, this is denoted by Equation 1:

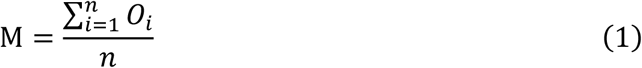

where, *n* is the number of recording channels and *O*_i_ denotes the original signal of the *ith* channel.

A simulation of how the averaging works is illustrated in Figure 2. Figure 2 (A) shows 5 synthetic signals randomly picked from a total of 18 signals that were produced to match the number of channel signals in the current dataset. Each signal was synthesised with: 1) a sine wave with a fixed phase across the 18 channels, with random amplitude, to simulate the mains noise signal; 2) a sine wave with random phase and random amplitude across the 18 channels, to simulate the EEG. Figure 2 (B) shows the separate components of the synthesized signals. Figure 2 (C) shows the noise and EEG components after being averaged across the 18 channels. As shown in Figure 2 (C), the simulated EEG signals had been averaged out while the simulated mains noise signal remained.

**Figure 2.**
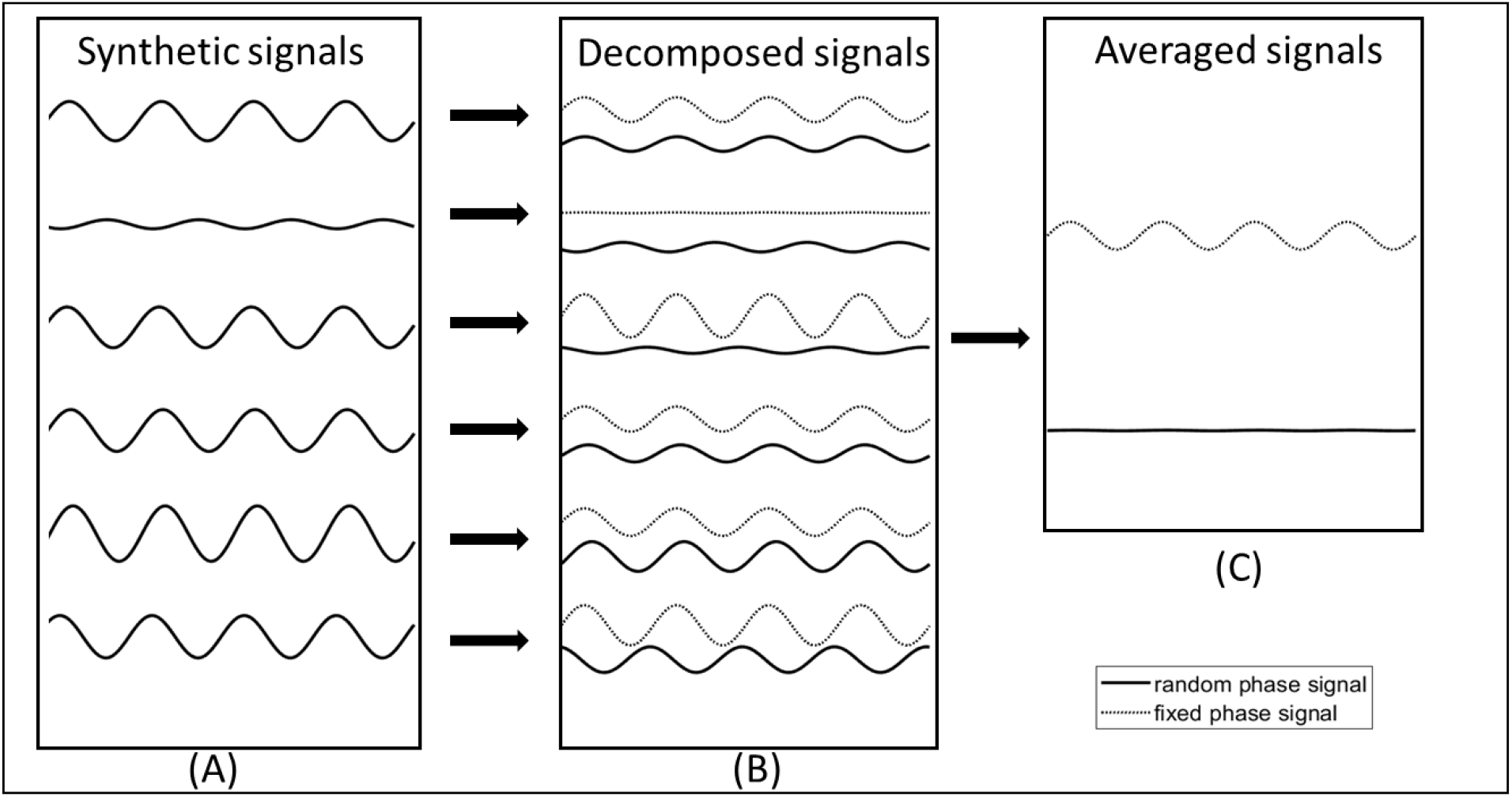
A set of simulated signals. Panel (A) shows 5 synthetic signals which were randomly picked from 18channels. Panel (B) shows two components of each signal, the dotted curve represents the phase fixed component, and the black solid curve represents the random phase component. Panel (C) shows the averaged of all (18) phase fixed signal and random phase signal separately. The result of averaging phase fixed signals is close to a flat line which means the simulated real brain components were eliminated, whereas the result of averaging random phase signals is a sine wave which means the simulated mains noise components were remained.

We simulated 1000 sets of such 18-channels EEG signal (each channel was generated by a sine wave with random phase and random amplitude) to test the influence of the number of channels. For each set of 18-channels EEG signal, we obtained 17 synthesized signals by averaging from 2 channels to 18 channels signal. Then the amplitudes of 17 synthesized signals were saved. This procedure was repeated 1000 times. The averaged amplitudes of 1000 sets of 17 synthesized signals were shown in Figure 3. We can see that as the number of channels increased, the amplitudes of synthesized signals are sharply decreased. It means that more channels will remove more background EEG signal and remain more pure mains power noise and bring more accurate results.

**Figure 3.**
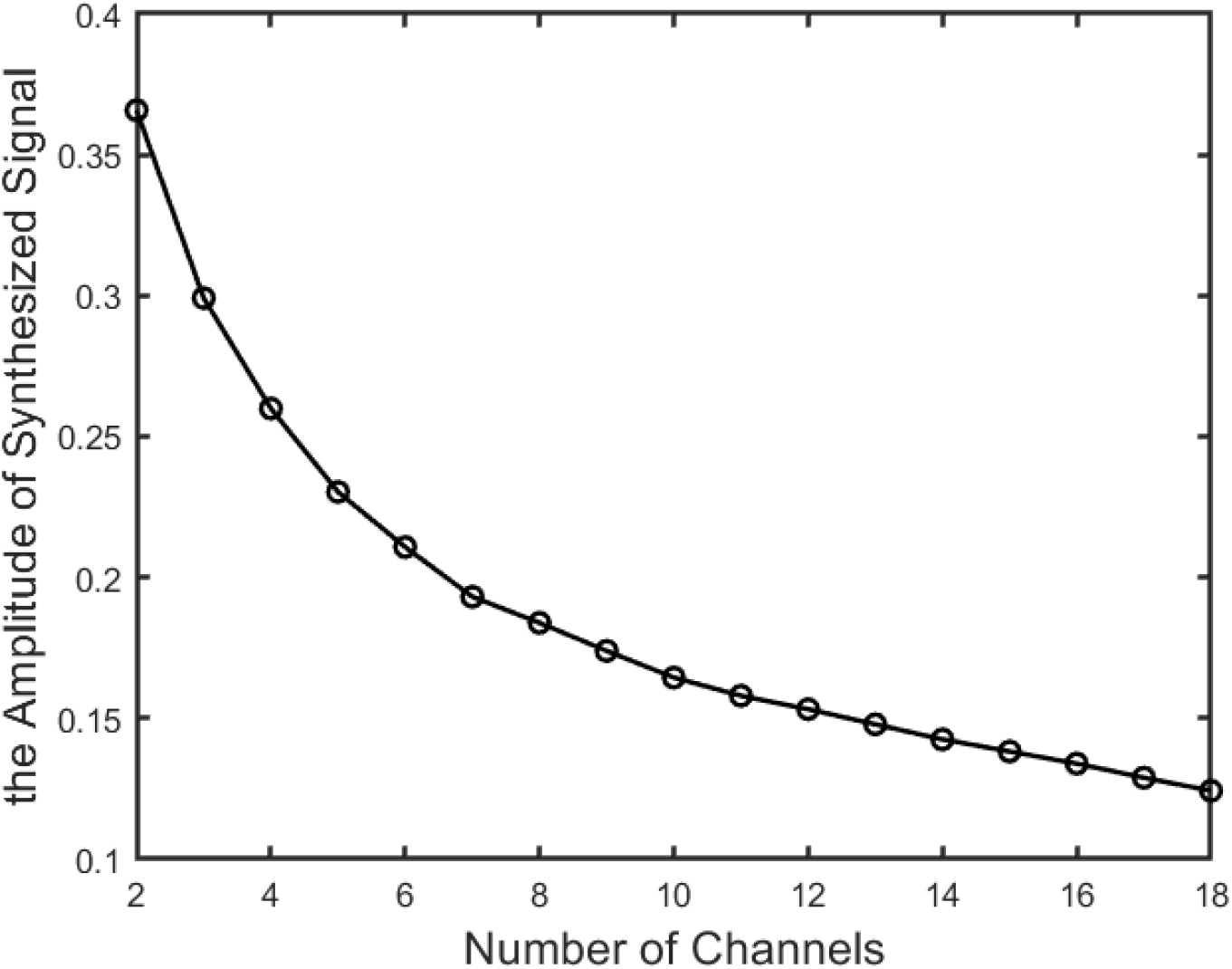
The influence of channel’s number. This figure shows the averaged amplitudes of 1000 sets of 17 synthesized signals. Each synthesized signal was obtained by averaging several (2-18) channels’ signal. We can see that as the number of channels increased, the amplitudes of synthesized signals are sharply decreased. It means that more channels will remove more background EEG signal and remain more pure mains power noise and bring more accurate results.

#### 2.4.3 Calculating the FFT phase of mains power noise frequencies

In the previous step, the approximate mains power signal (M) is obtained by Equation 1. In this step, we first apply a FFT algorithm to each signal, with a FFT window that is equal to the full length of the continuous signal, before any segmenting occurs. Matlab function fft() was used to compute the discrete Fourier transform (DFT) of *M.*

Next, we calculate the phase of signal M, for each frequency bin of the FFT output within the range of the mains noise. As a safeguard, the actual range is set slightly wider than the mains noise frequencies, e.g ~ 49.8 Hz to ~50.2 Hz. Matlab function phase() was used to compute the phases of each FFT frequency bin. Once the phases were obtained, they were used to generate sine waves to simulate the mains noise template (Equation 2).

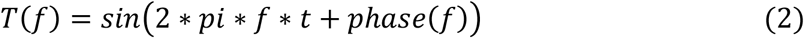

where, *f* is the frequency value which we are interested in; *t* represents time points.

#### 2.4.4 Calculating the FFT amplitudes of mains power frequencies

The mains power component propagates across channels retaining its phase while changing its size. Therefore, for each frequency value and channel, we scaled the template using the slope estimated by robust regression between *T* and the signal values. The scaled values were then subtracted from the residual EEG from previous iteration. The procedure was repeated for each frequency bin from ~49.8 Hz to ~50.2 Hz.

In the procedure, robust regression with a bisquare weight (Fox, 2002) function was used to scale the sine wave *T*() to each channel, because compared to ordinary least squares, robust regression can reduce the influence of outliers of the original signal.

#### 2.4.3 Iterative construction and removal of the mains power signal

After averaging, we calculate the phase and amplitude components of the mains noise signal. Once these are calculated, they are used to remove the mains noise signal. These steps are repeated for each frequency in a range that is slightly wider than the pre-determined mains noise frequencies, between ~ 49.8 Hz to ~50.2 Hz in this case.

The length of whole signal of our data samples is around 122880 and the sample rate is 256 Hz. Therefore, the frequency interval per sample is around 0.002 Hz, the band interval is 0.4Hz, and the number of iterations is around 200

## 3 Results

### 3.1 Removing mains power noise components

After the above steps, the mains power noise components of each channel were removed, and the natural EEG was left intact. Figure 1 shows a comparison of the raw signal and the signal after removal of mains noise in the frequency domain. The raw signal was randomly picked from one channel of one participant. Figure 1 shows clearly there is a mains power interference that exists in the range from 49.9 Hz to 50.1 Hz. After the mains power noise removal procedure, the mains power noise component was removed and only the natural EEG was left (Figure 4).

**Figure 4.**
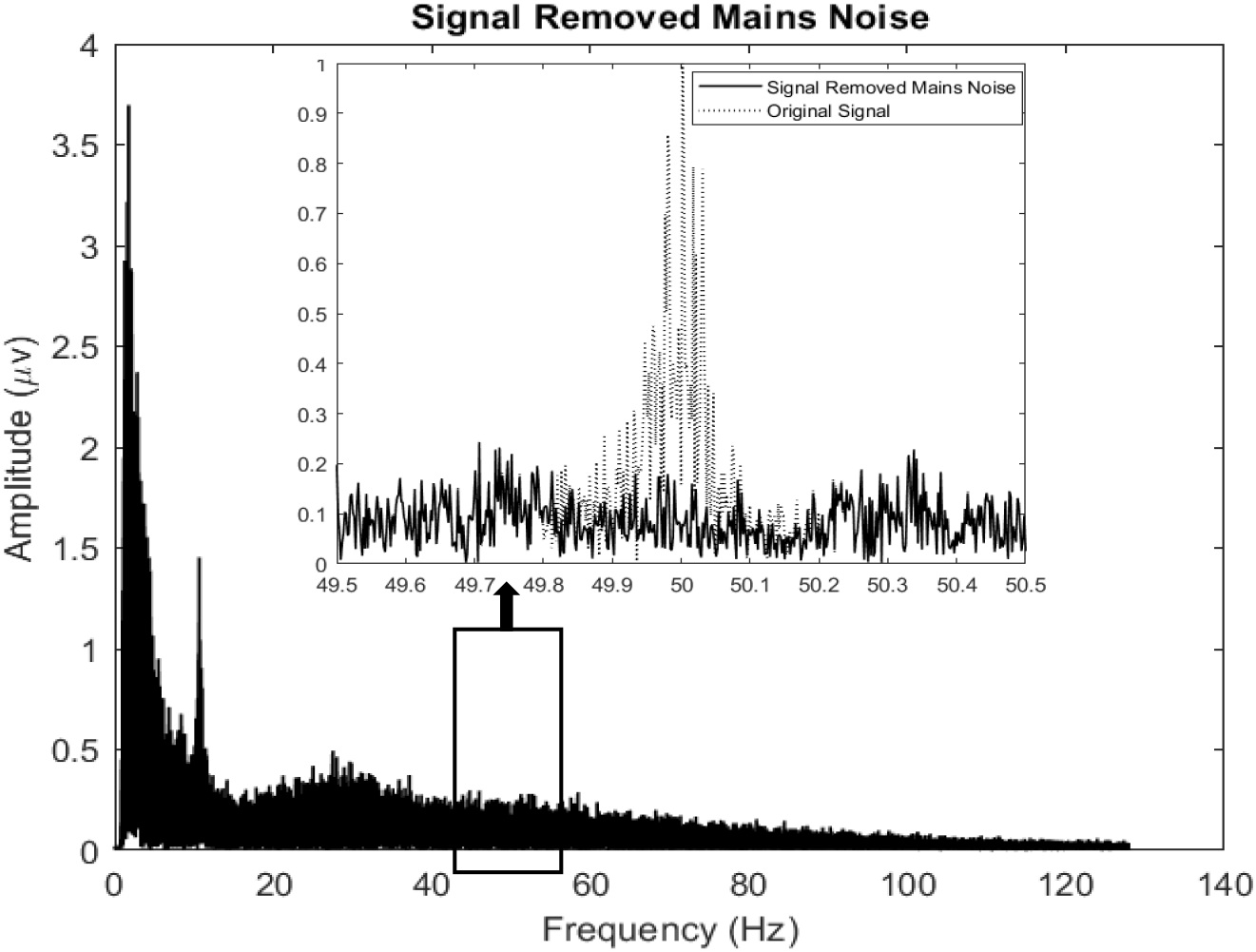
FFT spectrum after mains noise removal. Residual signal in Figure 1 after mains power removal with the proposed method.

### 3.2 Validation

#### 3.2.1 Mains noise simulation

To assess the effectiveness of our mains noise removal procedure, we added simulated noise waveforms to the existing EEG of all participants in our current dataset, and then tested the capacity of our mains noise removal method to recover the original EEG. We compared the results with common mains noise removal methods, ‘CleanLine’ and ‘multichannel PLI’.

To simulate mains noise, we reused the templates created above for noise removal (from 49.9 Hz to 50.1 Hz) for each channel. Each pair of phase and amplitude values of each mains power signal’s component were extracted and used to generate a sine wave to simulate a new mains noise component (Equation 3), with the frequency range 10 Hz shifted to 39.9 Hz to 40.1 Hz. These waves then sum together to simulate a mains noise signal.

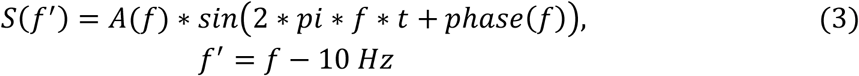

Where, *A*(*f*) and *phase*(*f*) are the pair of amplitude and phase values which were derived from original mains power signals. *S*(*f*′) is the new mains noise component.

The synthetic noise signal was added to the raw EEG. Figure 3A shows the data from a randomly selected participant in the time domain, with and without the addition of the mains noise signal. Figure 3B shows the same data in the frequency domain.

#### 3.2.2 Comparisons of our template method with CleanLine and multichannel PLI

As shown in Figure 5c-e, all the noise removal methods removed the simulated noise. However, CleanLine (Figure 5d) and multichannel PLI (Figure 5e) did not completely remove the simulated noise. To assess the degree of distortion on data pre-processed with stand steps, we split the original long signal into 2-s-length epochs with 1-s-length overlap and applied a Hanning window. Each epoch was then submitted to a FFT. The results were then averaged. Figure 5f shows the results. Compared to CleanLine and PLI, distortion to the original data was negligible with our method.

**Figure 5.**
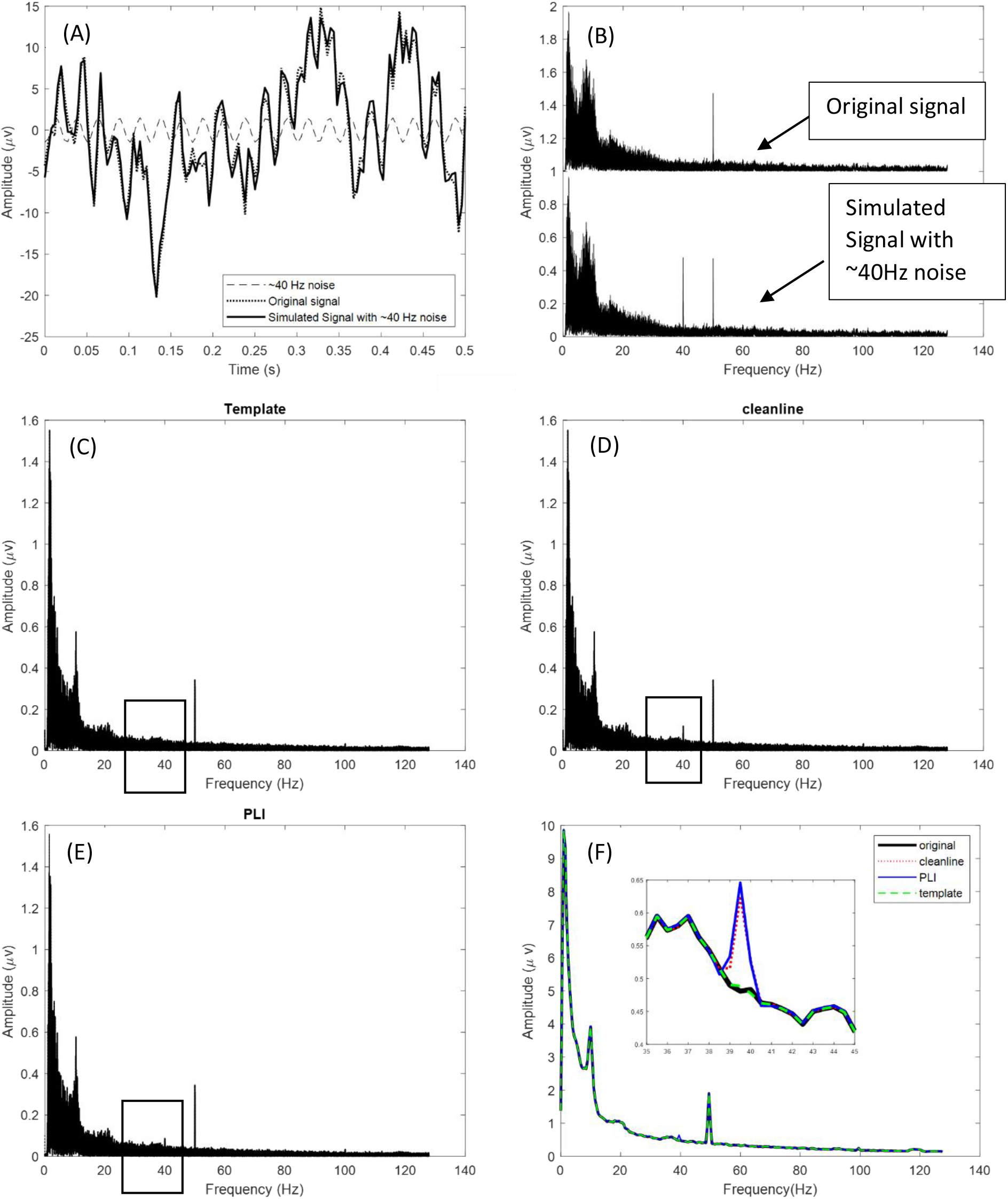
Synthesised mains power noise and testing. (A) Example of the simulated ~40Hz noise signal in the time domain from a randomly selected participant (B) The same data in (A) in the frequency domain. (C) Residual signal after our mains power removal procedure. (D) Residual signal after CleanLine noise removal. (E) Residual signal after multichannel PLI noise removal (F) Comparisons of the noise removal methods on data that were split into 2-s-length epochs with 1-s overlap, a Hanning window applied, followed by FFT and averaging.

To quantify the degree of fit of the noise-removed data to the original EEG, we calculated the correlation coefficient between the raw (i.e. before addition of simulated noise) and cleaned data for each method for all participants. Two sets of raw versus cleaned correlations were calculated for all participants. The first set was calculated using time domain signals and the other one was calculated in the frequency domain. Shared variances fo (r^2^ × 100) of the correlations were submitted to two one-way ANOVAs: (a) using the three sets of correlation values obtained by time domain signals of three methods; (b) using three sets of correlation values obtained by frequency domain signals of three methods. Figure 6 shows that the mean percent covariance is highest for the Template method in both time and frequency domain and lower for the CleanLine method. ANOVA including all correction methods found significant differences between the methods (time domain:F(2, 216) = 12.72, p < 0.001; Frequency domain: F(2, 216) = 17.44, p < 0.001). From Figure 6, we can see that mean correlation obtained by raw:cleaned signals of three methods are all above 99%. Thus all methods removed a substantial amount of the synthetic 40Hz mains power. However, the template method was significantly better than the other two.

**Figure 6.**
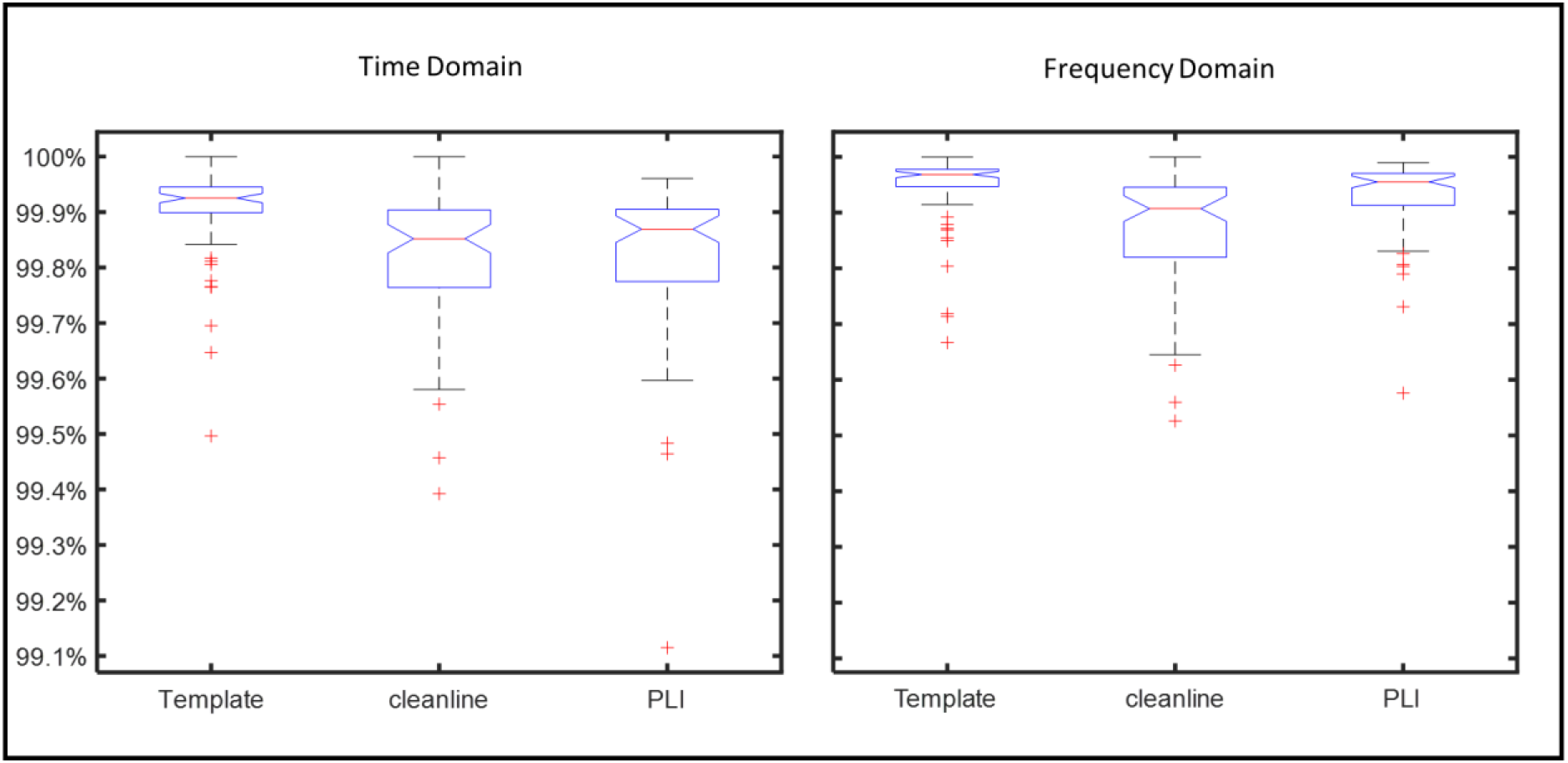
Percentage of covariance between the original and corrected signal in time/frequency domain for the different correction methods.

## DISCUSSION

EEG signals are contaminated by mains power from electrical equipment. The mains power causes a local peak in the EEG signal around the frequency of electricity. Here we proposed an automatic mains power noise removal method based on the physical basis that the mains power leads to the same phase components throughout all channels in the target frequencies. We have shown that this method can remove mains power noise components and with a success rate of over 99.9% in terms of recovering the variance of the original EEG prior to addition of synthesised mains power noise components.

Compared to Cleanline and PLI, which are commonly used for removing mains power noise artefacts (Brunner et al., 2013; Delorme and Makeig, 2004; Keshtkaran, 2014), our method produced the least distortion in both the time domain and frequency domain.

However, the current form of the model fails to 100% recover to original data, with minor distortion in the frequency domain. This effect is small,Compared to the current two common used methods (Cleanline, PLI). Note that the comparisons made here were based on 1Hz high passed filtered data. We did this for fair comparisons as Cleanline requires baseline drift removal for accurate electrical noise removal. Removal of the baseline drift is not actually a requirement of our proposed method. This makes it particularly suitable for studies on event-related potentials as these are known to be distorted by high pass filtering).

While our current method produces relatively undistorted data after denoising, its accuracy is proportional to information gain from the number of channels available.

